# The *Phlebotomus papatasi* transcriptomic response to trypanosomatid-contaminated blood is robust but non-specific

**DOI:** 10.1101/867382

**Authors:** Megan A. Sloan, Jovana Sadlova, Tereza Lestinova, Mandy J. Sanders, James A. Cotton, Petr Volf, Petros Ligoxygakis

## Abstract

Leishmaniasis, caused by parasites of the genus *Leishmania*, is a disease that effects up to 8 million people worldwide. Parasites are transmitted to human and animal hosts through the bite of an infected sand fly. Novel strategies for disease control, require a better understanding of the key step for transmission namely, the establishment of infection inside the fly. In this work we wanted to identify fly transcriptomic signatures associated with infection success or failure. We used next generation sequencing to describe the transcriptome of the sand fly *Phlebotomus papatasi* when fed with blood alone or with blood containing one of three trypanosomatids: *Leishmania major, Leishmania donovani* and *Herpetomonas muscarum*: a parasite not transmitted to humans. Of these, only *L. major* was able to successfully establish an infection in *P. papatasi*. However, the transcriptional signatures observed were not specific to success or failure of infection but a generalised response to the blood meal. This implies that sand flies perceive *Leishmania* as just a feature of their microbiome landscape and that any strategy to tackle transmission should focus on the response towards the blood meal rather than parasite establishment.

**Authors summary:** *Leishmania* are parasites that cause leishmaniasis, a group of serious diseases that affect millions of people, mainly across the subtropics and tropics. They are transmitted to humans by phlebotomine sand flies. However, despite establishment in the insect’s midgut being key to transmission, early infection events inside the insect are still unclear. Here, we study the gene expression response of the insect vector to a *Leishmania* parasite that is able to establish infection (*L.* major) one that is unable to do so (*L. donovani*) as well as one that is not a natural parasite of sand flies (*Herpetomonas muscarum*). We found that responses following any of the infected blood meals was very similar to uninfected blood meal. However, changes post-blood meal from day 1 to day 9 were dramatic. As a blood feeding insect can accumulate three times its weight in one blood meal, this seems to be the most important physiological change rather than the presence of the parasite. The latter might be just one in a number of microbes the insect encounters. This result will generate new thinking around the concept of stopping transmission by controlling the parasite inside the insect.

## Introduction

Leishmaniasis, a disease caused by parasites of the genus *Leishmania*, is endemic in 85 territories across the globe - with up to 8 million people affected^1^. Parasites infect vertebrates through the bite of an infected sand fly vector (Diptera: Phlebotominae). The acute form of disease, visceral leishmaniasis (VL) or kala-azar, is fatal in 95% of untreated cases and claims up to 50 thousand lives annually - though non-fatal infections causing dermatological symptoms are most common^1^. The ongoing VL elimination program in the Indian subcontinent is proving successful against the most severe clinical forms of VL^2^. However, elimination of leishmaniasis will likely require a combination of transmission blocking strategies and novel treatments. This is especially the case in light of reports of resistance to drugs used to treat human infections^3,4^, as well as pesticides used to control vector populations^5–7^. But to develop approaches to blocking transmission, we need a better understanding of the basic biology that underlines the interactions between parasite and insect vector.

The sand fly responses to blood feeding have been investigated with several gene families shown to be transcribed and/or expressed in response to a blood meal^8^. These include: digestive enzymes such as trypsins and chymotrypsins, pathogen recognition molecules and components of the peritrophic matrix – a protective chitinous mesh which lines the midgut after ingestion^8^. However, few sand fly genes or transcripts specifically associated with *Leishmania* infection. There is some evidence to suggest *Leishmania* are able to modify host responses to promote survival and infection establishment. Analysis of cDNAs isolated from dissected *Phlebotomus papatasi*^9^ and *Phlebotomus perniciosus*^10^ midguts revealed that several transcripts which are enriched after receipt of a blood meal are depleted when flies are fed blood containing *Leishmania*. These included digestive proteases, such as trypsins, as well as peritrophins which are chitin-binding components of the peritrophic matrix – a protective chitinous mesh which lines the midgut after ingestion^11^ and serves as a temporary barrier to leishmania^12^.

Recently, we described both the host^13^ and parasite^14^ transcriptomes in another trypanosomatid-insect infection model namely, *Drosophila melanogaster* and its own natural trypanosomatid parasite *Herpetomonas muscarum*. We showed that parasite feeding resulted in differential transcription of the two NF-κB pathways Toll and Imd, the dual oxidase pathway and STAT-dependant epithelial stem cell proliferation. Additionally, we found^14^ that the *H. muscarum* transcriptome during infection closely resembled that reported for *L. major* during *Phlebotomus duboscqi* infection^15^.

Given this, we wished to compare the *Drosophila* responses to those of sand flies during infection. Common transcriptomic signatures between the two systems would indicate an evolutionarily conserved response to trypanosomatid infection. Such responses would be of great interest for the development of broad-spectrum transmission blocking strategies for trypanosomatid diseases. However, no comparably comprehensive data’ is available from sand flies. As such we sought to describe the sand fly transcriptomic responses to trypanosomatid infection using next generation sequencing techniques (RNA-seq). Herein, we describe the transcriptome of *P. papatasi* at three timepoints corresponding to important stages of trypanosomatid infection; 1 day post blood meal (PBM); following blood meal digestion and when parasites can be found attached to the midgut epithelium (4 days PBM); when parasites have migrated anteriorly in the gut and are found in the thoracic midgut and the stomodeal valve of the fly (9 days PBM, Figure 1)^16^. Infections were done in the context of both permissive (*Leishmania major*) and refractory (*Leishmania donovani*) infections, as well as the with monoxenous (infects only insects) trypanosomatid *H. muscarum*, which is not a natural parasite of sand flies. Using this strategy, we hoped to identify host transcriptional signatures associated with permissive and refractory infection outcomes – in addition to identifying evolutionarily conserved host responses as described above.

**Figure 1.**
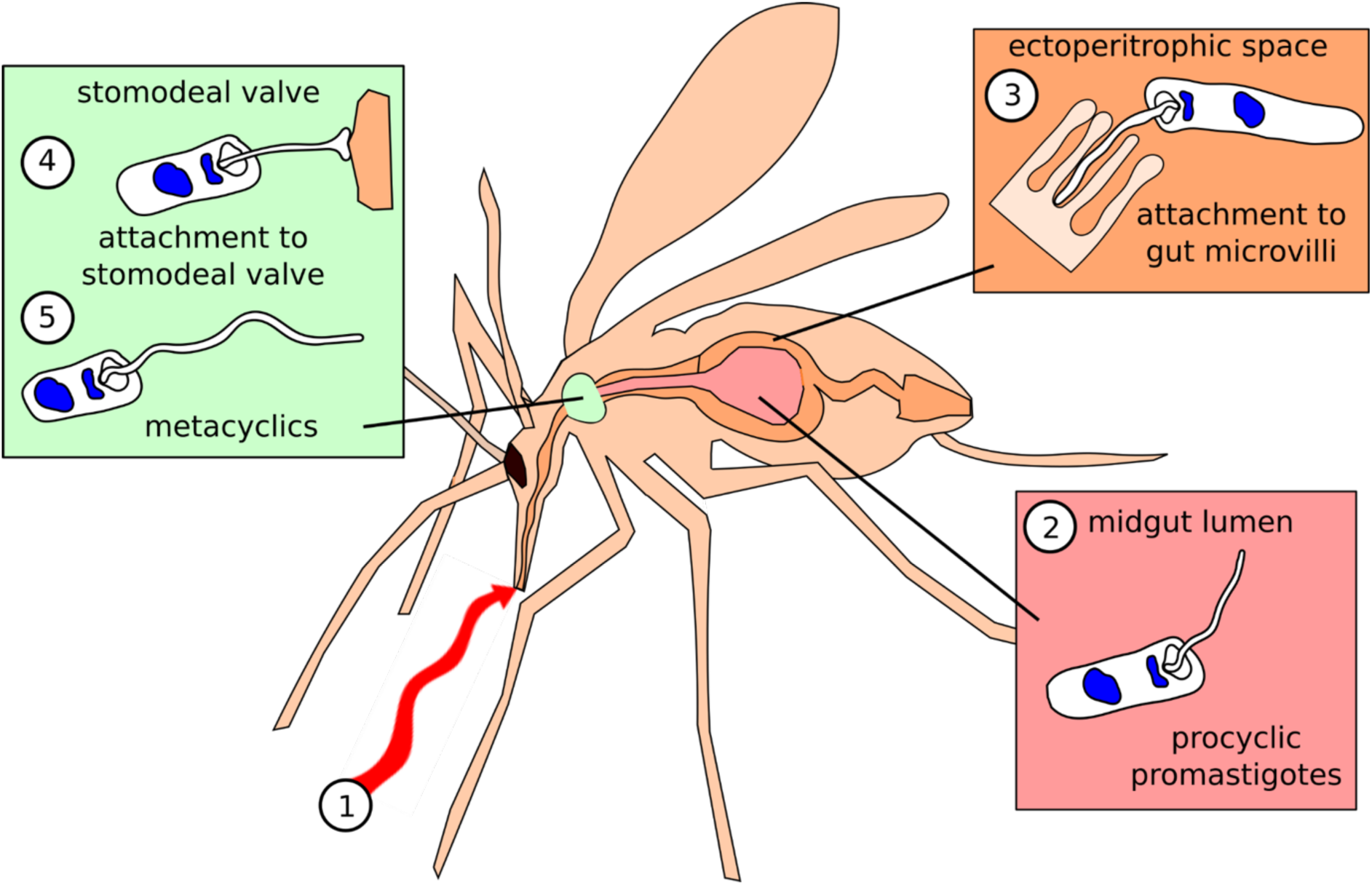
Schematic of the 3 major *Leishmania* stages in sand flies. Shortly after ingestion (**1**) of the blood meal promastigotes are localized in the midgut lumen, in the bloodmeal bolus surrounded by peritrophic matrix (**2**, 1 day post blood meal (PBM)). *Leishmania* wait until the PM is broken at the end of digestion, they enter the endoperitrophic space and attach to the epithelial wall (**3**) (> 4 days PBM). Finally, where parasites have migrated anteriorly to the thoracic midgut and the stomodeal valve of the fly and the human-infective metacyclic forms differentiate from the earlier stages (**4/5**, > 9 days PBM).

Our results indicate that whilst the sand flies’ transcriptomic response to blood feeding is robust and extensive, with the differentially regulation of thousands of genes – there is very little difference between the transcriptomes of blood fed and trypanosomatid (in blood) fed flies. Blood alone appears to trigger transcription of genes from several immune pathways – including Imd, Toll and JAK-STAT. Activation of these responses despite the absence of parasites in the meal may be a pro-active strategy by the sand flies to prevent infection.

## Results and Discussion

### Read mapping, *de novo* transcript assembly and differential expression analysis

RNA was purified from sand flies at 1, 4 and 9 days PBM and the resulting reads sequenced and mapped against the *P. papatasi* genome (Ppapl1, vectorbase^17^). The number of reads generated per sample ranged from 1.08-12.05 million reads with 69.7-79.3% mapping to the *P. papatasi* genome in each sample (Table S1). Upon visual inspection of read mapping using IGV^18^ it appeared that over 20% of reads were mapping to regions which lacked annotated features. To include these potentially novel genes in our analysis we assembled *de novo* concordantly mapped read pairs (from all samples) into 16,025 transcripts. The assembled transcripts were then merged with the existing annotation of 11,834 transcripts, to give a final set of 18,592 unique transcripts (see Supplementary data files). This represents approximately 97.2Mb of *P. papatasi* transcriptome with an average transcript length of 4,190 bp. All reads were then counted against the final set of transcripts for differential expression analysis.

Principal component analysis (PCA) showed a high degree of difference between the fly transcriptomes at day 1 PBM and those at day 4 or 9 PBM (Figure 2) – with transcriptomes from days 4 and 9 PBM appearing similar. We also note that samples do not clearly group in accordance with trypanosomatid feeding status.

**Figure 2.**
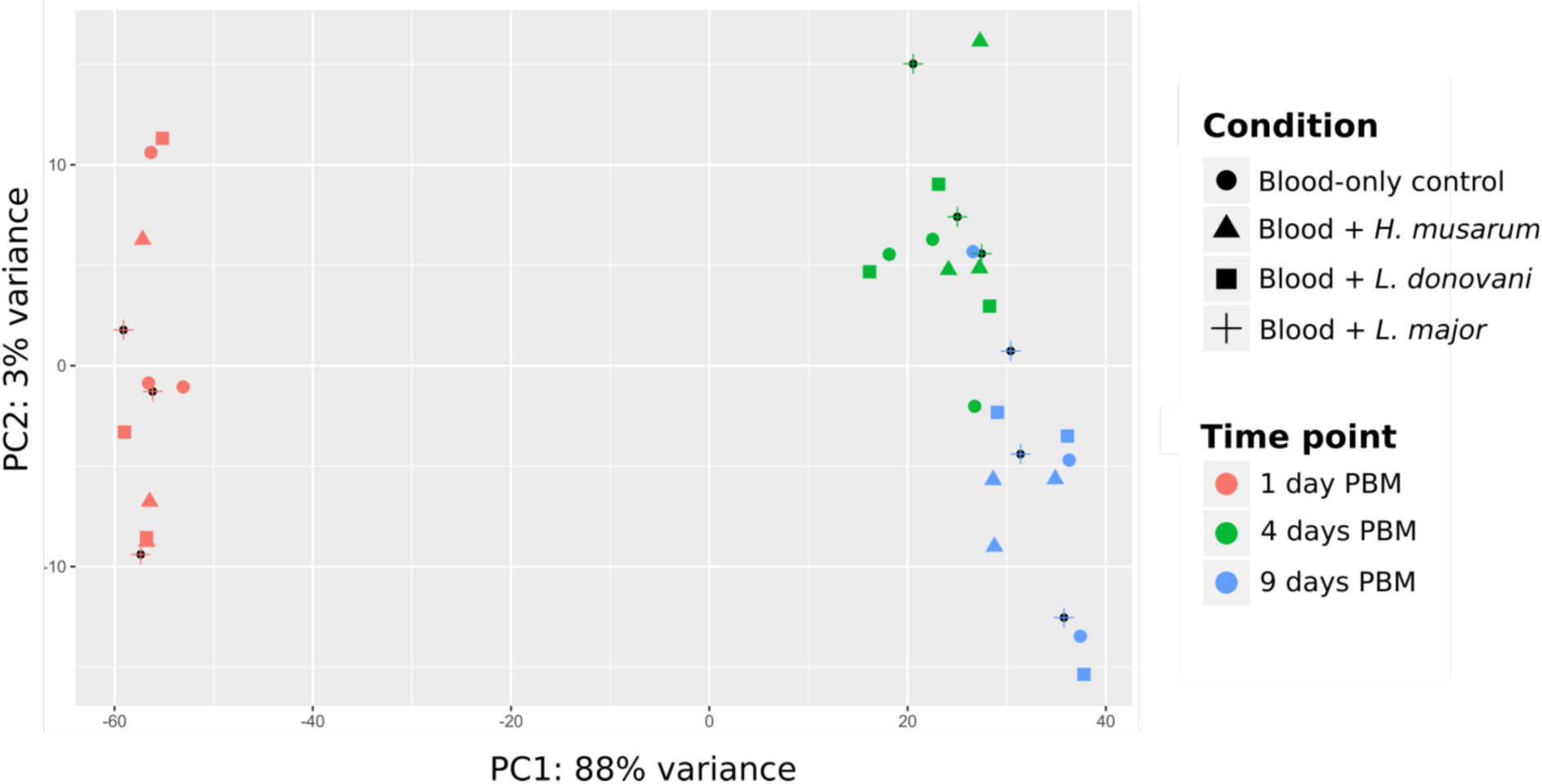
Principal component analysis shows that time was the major source of variation, not infection status (condition).

### Differential expression associated with trypanosomatid presence in the bloodmeal

There are few differentially expressed transcripts which were specifically associated with trypanosomatids being present in the blood meal (Table S2). We found no significant difference in transcript abundances between blood-fed and *L. major* fed flies at any time point. Furthermore, we find in excess of twelve thousand genes for which we reject the hypotheses that expression has changed by 2-fold or more in either direction in pairwise comparisons (Wald test) between blood only fed and trypanosomatid fed flies (Table 1).

**Table 1.** Numbers of transcripts which do not significantly change in expression by 2-fold or more in either direction between blood fed and trypanosomatid fed P. papatasi (p<0.05).

| vs. | Number of genes which do not significantly change in expression by 2-fold or more in either direction (p<0.05) |  |  |  |  |  |  |  |  |
| --- | --- | --- | --- | --- | --- | --- | --- | --- | --- |
|  | *Leishmania major* fed flies |  |  | *Leishmania donovani* fed flies |  |  | *Herpetomonas muscarum* fed flies |  |  |
|  | 1 day PBM | 4 days PBM | 9 days PBM | 1 day PBM | 4 days PBM | 9 days PBM | 1 day PBM | 4 days PBM | 9 days PBM |
| Blood fed 1 day PBM | 12586 |  |  | 12797 |  |  | 11957 |  |  |
| Blood fed 4 days PBM |  | 12188 |  |  | 12597 |  |  | 12356 |  |
| Blood fed 9 days PBM |  |  | 12762 |  |  | 12731 |  |  | 12634 |

We did however observe differential abundance for some transcripts after *H. muscarum* and *L. donovani* feeding compared to blood only fed control flies. There were significantly fewer transcripts for the gene PPAI009043, an orthologue to the *D. melanogaster* signalling protein Rho GTPase activating protein at 54D (RhoGAP54D), in flies fed *H. muscarum* than in blood fed controls at day 1 PBM (log2foldchange 1.13, p-adj < 0.05). The *Aedes aegypti* and *Anopheles gambiae* RhoGAP54D orthologues are upregulated in blood fed mosquitos compared to sugar fed controls^19,20^. Given this, and that this response was not seen after *Leishmania* feeding, this transcriptomic response may be *H. muscarum* specific. The biological significance of reduced RhoGAP54D transcription in this context remains unclear, however the protein is linked to epithelial morphogenesis during *Drosophila* development^21^ and so may also play a role in the mature insect gut.

In *L. donovani* fed flies there were significantly fewer transcripts for the putative transporter TrpA1 (PPAI004036, log2foldchange 2.8, padj < 0.05) versus blood only fed flies at 9 days PBM. TrpA1 is more generally associated chemo- and thermo-sensing^22,23^ in *Drosophila*, however a study by Du *et al*. 2006 links TrpA1 to the expulsion of food-borne pathogens by increased defecation and the DUOX pathway (discussed further below)^24^. Speculatively, reduction in TrpA1 transcripts after *L. donovani* feeding may hint at modification of host defensive pathways to promote survival. We also find significantly more CUFF.12679 transcripts (log2-foldchange 16.8, p-adj < 0.05) in *L. donovani* fed flies. This novel transcript lacks conserved domains or sequence similarity to known Dipteran gene transcripts.

Direct comparisons between trypanosomatid infections yielded similarly few differentially expressed transcripts (Table S3). At day 1 PBM the only differentially expressed transcript between the three infections was that of trypsin 1 (PPAI010956, padj < 0.05) which was 2-fold enriched in *H. muscarum* fed flies compared to those fed *L. donovani*.

After defecation at around 4 days PBM it is thought only parasites able to establish in the ectoperitrophic space persist to develop mature infection^25^. Despite the differences in the infection outcome reported in laboratory infections across the three trypanosomatids^16^, there were few differences in the host transcriptome at this critical time point. Two transcripts were found to be significantly differentially abundant - one corresponding to the PPAI000999 gene and the other a novel transcript CUFF.14170. Both transcripts were found at significantly higher levels (p-adjusted <0.05, log2foldchanges of 4 and 18 respectively) in *H. muscarum* fed flies compared to those fed *L. donovani*. PPAI000999 encodes a protein predicted to bind to chitin (GO:0006030, GO:0008061 and smart00494). The novel transcript CUFF.14170 has no known conserved domains and BLAST searches against Dipteran sequences did not yield any significant hits.

The most variation between the three infections was found at 9 days PBM, where 6 transcripts were differentially expressed between *Leishmania* fed and *H. muscarum* fed flies (padj < 0.05). Flies fed *L. donovani* had significantly more transcripts for the previously discussed TrpA1 (PPAI004036), and significantly less for the putative zinc metalloprotease PPAI010164 and novel transcript CUFF.12679, than those fed *H. muscarum*. Flies fed *L. major* had significantly more transcripts for the hypothetical protein PPAI002947. Additionally, feeding with *H. muscarum* resulted in significantly more CUFF.14170 transcripts, a novel transcript from this study which lacks conserved domains, than both *Leishmania* infections (padj < 0.05).

Overall, the above observations suggest that blood feeding status is the major source of transcriptional variation in these flies and not trypanosomatid infection. As such we further investigated transcriptomic changes after blood feeding alone in *P. papatasi*.

### The *Phlebotomus papatasi* transcriptome after blood feeding

Ingestion of blood alone was associated with significant changes to the transcriptome. The transcriptomes at day 1 PBM appeared very different to those at 4 (and 9) days PBM, with 12,289 significantly differentially regulated transcripts (Table S4). However, after defecation of the blood meal remnants the transcriptome was comparatively stable with 264 differentially regulated transcripts (4 vs. 9 days PBM, Table S5). Due to the large number of differentially expressed transcripts highlighted by these comparisons we first investigated transcripts whose log2 fold change was > 4 in either direction between timepoints. From this subset we were able to focus our analysis on a number of key genes and pathways which are discussed further below (Tables S6 and S7).

### Early transcriptomic responses to blood meal ingestion are concerned with digestion, metabolism and immunity

Of the 217 transcripts differentially regulated > 4-fold between 1 and 4 days PBM, 197 transcripts were found to be comparatively enriched at day 1 PBM and 20 were comparatively enriched at day 4 PBM. Ninety-eight of these transcripts did not contain known conserved domains.

Transcripts for putative and known trypsins were one of the most highly represented groups differentially regulated between day 1 and day 4 PBM. We observed upregulation of 9 transcripts for putative trypsins and chymotrypsins – including the previously characterised chymotrypsins 1 (PPAI010833), chymotrypsin 3 (PPAI005023) and trypsin 4 (PPAI010456)^8,11,26^. We also observed upregulation of transcripts which may represent novel trypsins, based on conserved domains and similarity to other Dipteran trypsin/chymotrypsin sequences, as they are not included in the current genome annotation (Ppap v1^17^) (CUFF.11666, CUFF.9493, CUFF.6542) and chymotrypsins (CUFF.15058, CUFF.16005, CUFF.15086, CUFF.14587, CUFF.12454). In contrast, the transcript putatively encoding for trypsin 1 (PPAI010956) was shown to be enriched at day 4 PBM compared to the earlier timepoint. The roles of trypsin and chymotrypsin-like serine proteases during blood digestion in hematophagous insects are well characterised with expression levels varying according to type and the time since the last blood meal. Our findings agree with previous work, which showed upregulation of trypsins 3/4 and chymotrypsin 1 in response to the blood meal, as well as the decrease of trypsin 1^26^.

In addition to the trypsins themselves, five transcripts whose products are predicted to contain trypsin inhibitor like domains (PPAI003932, PPAI000270, PPAI000272, PPAI000274, PPAI003557) were also comparatively enriched at day 1 PBM (vs. day 4 PBM). It is possible the corresponding proteins play roles in the regulation of the trypsin 1 as well as other trypsins (e.g. trypsin 2), reported to be downregulated after blood feeding^26^.

Several transcripts encoding for proteins with predicted serine protease/proteolytic activity, the sequences of which do not resemble trypsins/chymotrypsins were also comparatively enriched at day 1 PBM. These included two known genes (PPAI009419, PPAI009871) and three novel transcripts (CUFF.6132, CUFF.6133, CUFF.16132). Serine proteases are implicated in several other cellular processes including innate immune signalling – notably in Toll pathway activation^27^ and the melanisation response^28^. The predicted protein for PPAI009419 shares approx. 51% identity with the *Culex quinquefasciatus* CLIPA15 (also known as masquerade) across its sequence. CLIPA proteases interact with and regulate other CLIPs, and the prophenoloxidases (PPO), involved in the melanisation repsonse^29,30^. This response produces reactive quinones which then polymerise to form the dark insoluble pigment melanin. These molecules can encapsulate and isolate invading pathogens or toxic compounds. They also locally generate high local levels of cytotoxic reactive oxygen species and prevents gas diffusion, starving the invading pathogen of oxygen. In addition to the putative CLIPA transcript, four pro-phenoloxidase transcripts are upregulated early infection (PPO1 - PPAI008831, PPAI010450; PPO2 - PPAI012836, PPAI012835). These zymogens are the rate limiting enzymes in the production of melanin. PPO1/2 and CLIPA15 were also upregulated immediately after blood feeding in *Anopheles gambiae*^20^ - suggesting this is a conserved response to blood feeding in Dipterans.

We also observed differential transcription of another group of proteins reported to play vital roles in protection against invading pathogens - peritrophins. These core components of the peritrophic matrix (PM) have been shown to be a major barrier against infection establishment. Knockdown of Peritrophin 1 (Per1) in *P. papatasi* results in an approximately 40% increase in *Leishmania major* load at 48 hours after parasite ingestion^31^. In our study, Per1 transcripts were highly enriched at day 1 PBM (vs. day 4 PBM) with log2 fold change of 9.96. Of the 32 annotated peritrophins in the *P. papatasi* genome, 14 were found to be significantly differentially regulated between days 1 and 4 PBM (Table 2). The majority of transcripts were comparatively enriched at day 1 PBM, however Per2 and Per28 transcripts were more abundant at later timepoints. Ramalho-Ortigão *et al*. 2007^10^ showed that *P. papatasi* peritrophin 1 (Per1) transcripts were enriched in flies fed a blood meal compared to a sugar meal, whilst peritrophin 2 (Per2) transcripts were comparatively depleted in blood fed flies. Additionally, the group showed that transcripts for both Per1 and Per2 were depleted in *L. major* infected flies compared to those fed only blood^11^. Our data largely agree with these findings. However, transcript levels were not statistically significant different between trypanosomatid and blood-only fed flies - though we do observe fewer transcripts for Per2 (PPAI009723) in trypanosomatid-fed flies at day 4 PBM (Figure S1). Other than Per2, the patterns in peritrophin transcript abundance for trypanosomatid fed flies resembled those of the blood fed controls.

**Table 2.**
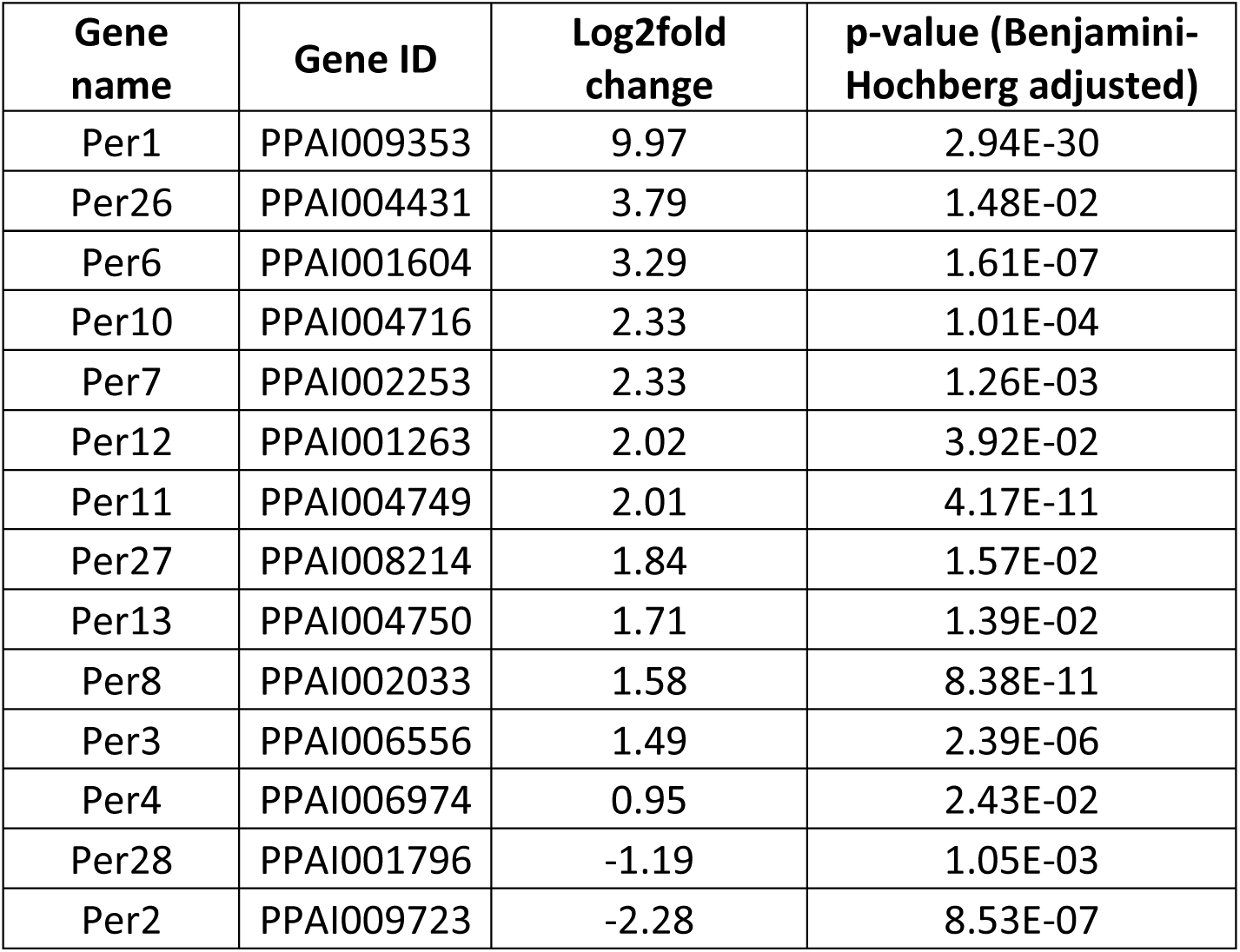
*P. papatasi* peritrophins significantly differentially regulated between 1 and 4 days PBM. Positive fold change values indicate enrichment at 1 day PBM and negative values indicate enrichment at day 4.

Additionally, transcripts for another chitin-binding protein, PPAI000188, were significantly more abundant at 4 days PBM than at day 1 PBM. The sequence of PPAI000188 resembles the *Lutzomyia longipalpis* protein ChiBi (EU124616.1^32^, 84% protein sequence identity). ChiBi was shown to be enriched *in L. longipalpis* fed with blood containing *L. infantum chagasi*^32^. Its upregulation here *in P. papatasi* in the absence of *Leishmania* may indicate this upregulation is a more general response to blood meal, rather than an infection-specific response.

In addition to trypsins, transcripts of several other groups of genes associated with digestion and nutrient uptake were differentially regulated PBM. Several transcripts for lipid metabolism associated genes were found to be upregulated at day 1 PBM. In addition, eight transcripts corresponded to known extracellular carboxylic ester hydrolases (PPAI002323, PPAI003061, PPAI003086, PPAI005115, PPAI005116, PPAI005680, PPAI009133, PPAI008993). Similarly, transcripts for a putative sterol transfer protein (PPAI008838), and two paralogous membrane fatty acid desaturase genes (PPAI008098 and PPAI002108) were shown to be comparatively enriched at day 1 PBM. One transcript, CUFF.7417, does not correspond to a known gene, however the transcript showed strong sequence similarity to the extracellular carboxylic ester hydrolases paralogues PPAI005115 and PPAI005116 mentioned above (90% identity). Additionally, CUFF.7417 is immediately downstream of PPAI005115/6 in the genome and as such we propose this represents a previously unknown paralogue.

Four transcripts coded for proteins with solute carrier domains (cl00456). These transcripts encode for the two paralogous sodium-coupled monocarboxylate transporters (SCMTs, PPAI005125 and PPAI007402) and two putative SCMTs (CUFF.14648 and CUFF.14649). The SCMTs are transmembrane proteins, which move molecules with a single carboxylate group including pyruvate and lactate, across the plasma membrane in a proton-dependent manner and are associated with the insect midgut brush boarder^33^.

We found two transcripts, CUFF.17209 and CUFF.15972, whose products are predicted to contain the conserved insect allergen related repeat domain (pfam06757). These transcript sequences also showed similarity to reported cDNAs for *P. papatasi* microvillar proteins MVP1 and 2 respectively (>89% identity to mRNA sequences). These proteins were also found previously to be upregulated in sand flies upon ingestion of a blood meal compared to sucrose-fed flies^11^. These transcripts could not be assigned to an annotated gene in the current vector base genome (Ppal1^17^). The function of these proteins is not well understood though they appear to have a conserved signal peptide at the n-terminus and lack transmembrane domains.

Finally, three olfactory (Or57 - PPAI013155, Or99-PPAI013290 and the putative protein PPAI002404) and a gustatory receptor orthologous to sweet taste receptors of *Drosophila* (Gr9 - PPAI010978), were upregulated at day 1 PBM compared to later timepoints. It is likely these sensory receptors were involved in sensing and acquisition of the blood meal and subsequent decreases in their transcript abundances, may indicate these sensors were not required after digestion.

### The transcriptome after defecation of the blood meal is comparatively stable

The two later timepoints in this study had similar transcriptomic signatures, with only six transcripts comparatively enriched >2-fold at 9 days PBM (vs. 4 days). These transcripts corresponded to two glutamate receptors (PPAI003634, PPAI008275), apoptosis inhibitor survivin (PPAI002284), two histone methyltransferases (PPAI005539, PPAI005538) and a mucin (PPAI009152). Mucins have been implicated in the interaction with *Leishmania* parasites. Given that several immunity-related transcripts (including peritrophins, mucins and melanization pathway genes) were upregulated, we postulated that upon blood meal ingestion a general immune response was triggered. As such we investigated the transcription of the members of the two major innate pathways after a bloodmeal: Toll and Immunodeficiency (Imd). Both pathways have been shown to play a role in the response to trypanosomatids^13,34–38^. Furthermore, we also investigated members of the Dual-oxidase (DUOX) and JAK-STAT pathways, both of which were implicated in *D. melanogaster*-*H. muscarum* interaction^13^. Differential regulation statistics for these transcripts can be found in Table S8.

### Blood ingestion alone is associated with increased innate immune gene transcription

In blood-fed flies, transcripts putatively encoding early Toll pathway genes (two Toll receptors, Spätzle and GNBP3) were found to be significantly enriched at day 1 PBM compared to days 4/9 PBM (fold change > 2, p-adjusted < 0.05, Figure 3A). An exception to this was the *spätzle processing enzyme* (SPE) the putative transcript for which is enriched in the latter two timepoints along with several intracellular Toll pathway components. These trends were broadly consistent in blood-only fed flies as well as those fed with each of the trypanosomatids. However, only flies fed with blood containing *L. major* or *L. donovani* promastigotes had significantly higher levels of transcripts encoding Toll pathway inhibitor Cactus at day-1 PBM compared to day 4 PBM (> 2-fold, p-adj < 0.05). Cactus transcript abundance was not significantly different between days 1 and 4 PBM in blood only or *H. muscarum* fed flies.

**Figure 3.**
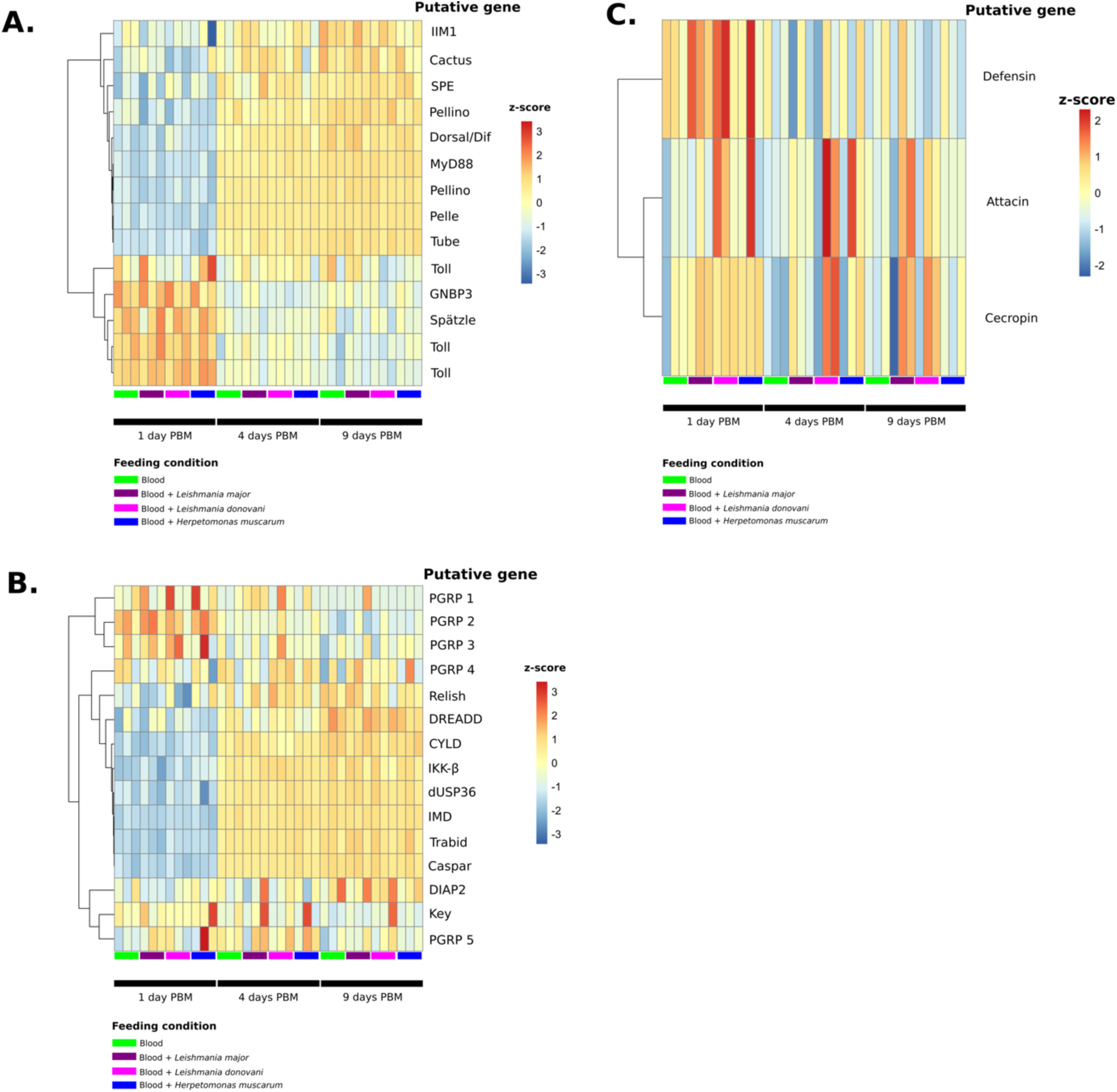
Transcription of genes form the two major innate immune pathways in *P. papatasi* across samples. **A.** A heatmap of z-scores (based on log transformed, normalised counts data) for Toll pathway genes across samples. **B.** A heatmap of z-scores (based on log transformed, normalised counts data) for Imd pathway genes across samples. **C.** A heatmap of z-scores (based on log transformed, normalised counts data) for anti-microbial peptide genes across samples.

A similar pattern emerges for the IMD pathway (Figure 3B). Transcripts for putative peptidoglycan recognition proteins (PGRPs) were more prevalent at day 1 PBM compared to later timepoints (Figure 6B). However, only putative PGRP 2 (CUFF.5670) was found to be statistically significantly enriched (2.23-fold) at day 1 PBM (vs. day 4/9 PBM, p-adj < 0.05). The transcripts putatively encoding IMD, and several other proteins downstream of IMD in the pathway were found to be significantly enriched (padj < 0.05) at 4 and 9 days PBM (vs. day 1 PBM), including: DREDD, TAK1 and IKKβ. We also observed significant enrichment of transcripts putatively encoding negative regulators of the IMD pathway Caspar, dUSP36, Trabid at days 4 and 9 PBM (>2 fold, p-adj < 0.05). Interestingly, the IMD transcription factor Relish was not significantly differentially regulated in blood-only fed flies, however flies fed blood containing *L. major* or *L. donovani* promastigotes showed enrichment of putative Relish transcripts at day 1 PBM compared to at 4 and 9 days PBM. As such, whilst there is overall upregulation of IMD pathway transcription with or without trypanosomatids in the blood meal, there may be important differences in the expression levels of the innate effectors the meal regulates when *Leishmania* are present.

Both Toll and IMD result in the expression of a suite of antimicrobial peptides. Transcripts for these immune effectors were not significantly differentially regulated after blood feeding alone. However, flies fed with blood containing *L. major* or *L. donovani* promastigotes were found to have significantly more transcripts for the AMP defensin at day 1 PBM compared to day 4 PBM (2.3 and 1.75 respective log2foldchanges, padj < 0.05). This was not observed in *H. muscarum* fed flies. Whilst transcript levels for other antimicrobial peptides did change after trypanosomatid feeding, e.g. we observe elevated transcript levels for cecropin and attacin in some trypanosomatid infections (Figure 3C), overall these were not found to be statistically significant changes.

In addition to AMP expression, the IMD pathway can also result in the transcription of the NADPH oxidase, dual-oxidase (DUOX), through interaction of IMD with MEKK1^39^. This transmembrane protein is responsible for production of ROS in the gut epithelium in response to microbes. We found that DUOX transcripts were significantly higher at days 4 and 9 PBM compared to day 1 PBM in all feeding conditions (log2foldchange 2.98-3.33, p-adj < 0.05, Figure 6C) – with no significant difference in DUOX transcript abundance between 4 and 9 days PBM in any infection condition. Similarly, we see significant increases in transcripts for genes upstream of DUOX across infection conditions including: the transcription factor ATF2, p38 kinase and MEKK1. As such, induction of DUOX pathway transcription appears to be a generalised response to blood feeding rather than an infection-specific response.

### The JAK-STAT pathway is also associated with the dipteran response to trypanosomatids

Finally, given the association between the JAK-STAT pathway (Figure 4B), dipteran gut morphology and immunity^40^, particularly in a trypanosomatid infection context^13^, we also investigate the transcription of key components of this pathway after blood feeding. We observed higher abundance of putative Upd1 transcripts at day 1 PBM compared to later timepoints, however this change was only show to be statistically significant for flies fed with blood and *L. major* where there was a 2.2-fold enrichment of putative Upd1 transcripts. Furthermore, putative transcripts for the JAK-STAT transcription factor STAT92E were 2-fold enriched in flies in all infection conditions at the two later time points (vs. 1 day PBM, p-adj < 0.05). We also observed a modest enrichment of transcripts for cytokine Upd2 and the transmembrane receptor Domeless at days 4 and 9 PBM compared to earlier timepoints (fold changes 1.19 and 1.74 respectively, p-adj < 0.05). The transcription pattern for signalling protein hopscotch resembled that of Domeless, however these transcripts were only found to be statistically significantly enriched in trypanosomatid fed flies (padj < 0.05). Together these observations suggested an increase in JAK-STAT signalling a few days after a blood meal in *P. papatasi*. Further work to investigate if this signalling translates to changes in gut homeostasis, such as the increased stem cell proliferation observed in the *Drosophila*-*Herpetomonas* model, will be important. Currently however, as transcript abundance for STAT92E is enriched in blood only fed controls this response does not appear to be trypanosomatid specific.

**Figure 4.**
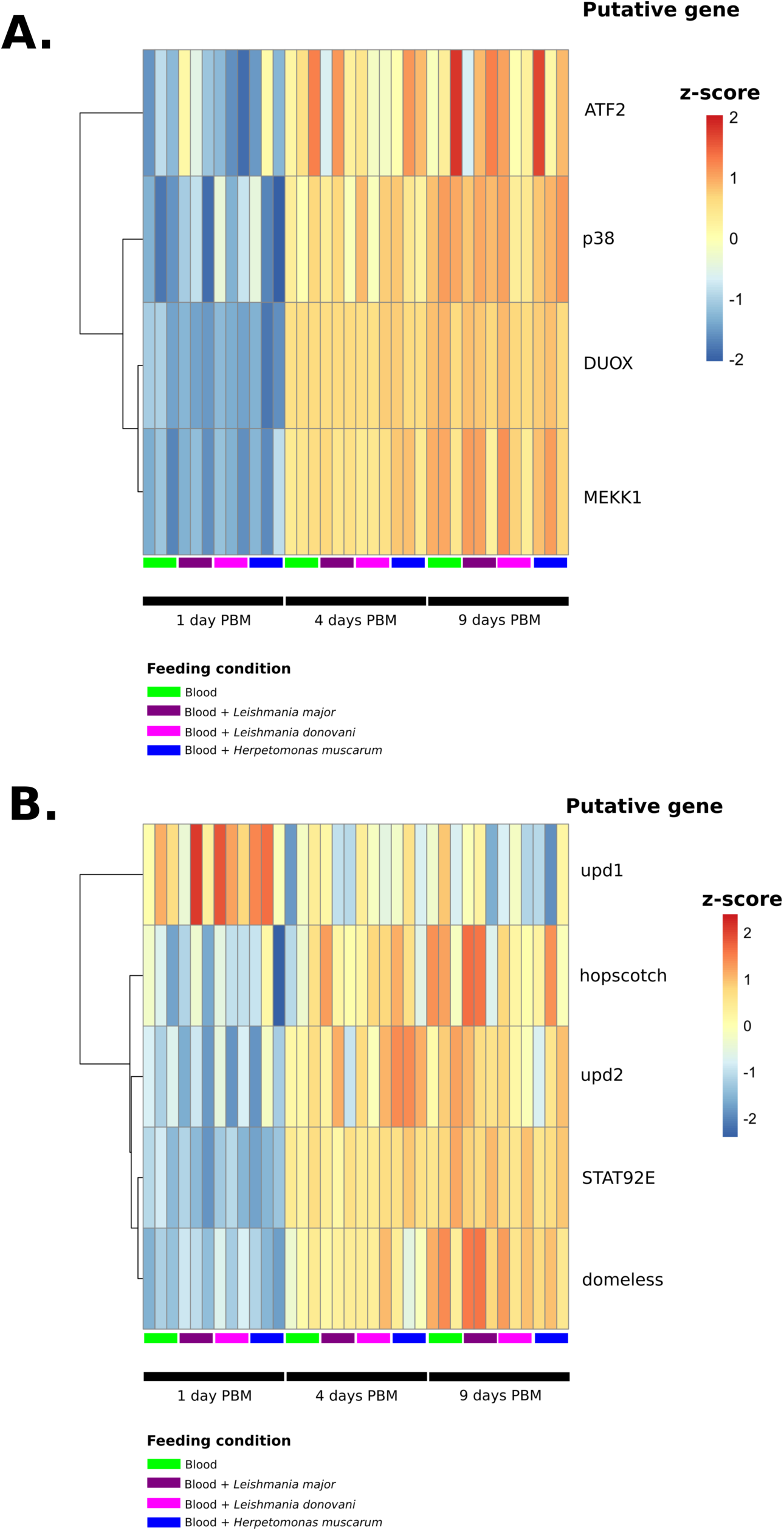
Transcription of genes form the DUOX (A) and JAK-STAT signalling pathways in *P. papatasi* across samples. Heatmaps of z-scores (based on log transformed, normalised counts data) across samples.

## Concluding remarks

Given the magnitude of the transcriptomic changes associated with blood feeding alone, and the little variation between blood meals spiked with trypanosomatids that produce very different infections we speculate that aforementioned defensive responses are not infection specific. Such a strong response to the blood meal alone is not surprising given the additional stresses associated with the hematophagous habit^41^. The high-risk nutrient attainment method drives the insects to take large volumes of blood at each meal e.g. mosquitoes and tsetse flies expand up to 3 times their pre-meal size a blood meal^41,42^ – putting enormous mechanical strain on the tissues. In addition to the volume, the content of their meal presents additional problems: excess water/ions^43^, toxic compounds^44^ and bacterial expansion in response to the rich meal^45,46^. Due to the warm-blooded nature of their victim’s blood temperature of blood-feeding arthropods can rapidly (< 60 seconds) increase by over 10°C during their meal^47,48^. All of which must be overcome even in meals which do not contain parasites. Additionally, activation of immune pathways in absence of known infection may be a strategy to ‘pre-emptively’ protect the host against pathogens/toxic compounds which may be present in the newly ingested meal. Such anticipatory responses have been reported in other hematophagous insects including malaria vector *Anopheles gambiae*^49^.

Moreover, it is known that in sand flies, the blood meal is followed by a decrease in overall gut bacterial diversity^50^ coupled to an increased abundance of aerobic bacteria^46^. We suppose that these changes may mask any effects from the presence of trypanosomatids. However, there was also no significant difference between day 9 PBM *L. major* and the other day 9 infections. This is important since by day 9, the blood meal has long been digested and it is only *L. major* that is left while the other parasites are cleared. This underlines the non-specificity of the *P. papatasi* response and implies that for the fly, *L. major* is just another feature in the microbiome landscape.

## Materials and methods

### *Phlebotomus papatasi* maintenance

A laboratory colony of *P. papatasi* (originating from Turkey) was maintained in the insectary of the Charles University in Prague under standard conditions (at 26°C fed on 50% sucrose, humidity in the insectary 60-70% with a 14 h light/10 h dark photoperiod) as described previously^51^.

### Trypanosomatid maintenance

*L. donovani* (MHOM/ET/2010/GR374), *L. major* LV561 (LRC-L137; MHOM/IL/1967/Jericho-II) and *H. muscarum*^13^ were cultured in M199 medium (Sigma) containing 10% heat-inactivated foetal bovine serum (FBS, Gibco) supplemented with 1% BME vitamins (Basal Medium Eagle, Sigma), 2% sterile urine, 250 μg/ml amikacin (Amikin, Bristol-Myers Squibb) at 23°C (*L. donovani, L. major*) or 28°C (*H. muscarum*).

### *Phlebotomus papatasi* infections

*Leishmania* and *H. muscarum* promastigotes from log-phase cultures (day 3-4 post inoculation) were resuspended in defibrinated and heat-inactivated rabbit blood (LabMediaServis) at concentration 1×10^6^ promastigotes per mL which corresponds to 500-1000 promastigotes per *P. papatasi* female ^25^. Sand fly females (5-9 days old) were infected by feeding through a chick-skin membrane (BIOPHARM, Czech Republic) on the suspension. Engorged sand flies were maintained in the same conditions as the colony. Females were dissected at days 1, 4 and 9 post bloodmeal (PBM).

### RNA extraction

RNA was extracted from pools of 10 sand flies from each condition and timepoint. Whole flies were homogenised in 200μL TRIzol Reagent (Thermofisher) before 300ul more TRIzol was added. The homogenate was mixed and incubated at 4°C for an hour. Debris was then spun down by centrifugation at 12, 000 ×g for 5 mins and the resulting supernatant transferred to a new tube. To each sample 100 μL of chloroform was added and samples incubated on ice for 2-3 minutes. The three phases (phenol-chloroform, interphase, and upper aqueous phase) were separated by centrifugation at 12, 000 ×g for 15 min at 4°C and the upper phase containing the RNA was moved to a new tube. Following this RNA extraction proceeded according to the TRIzol Reagent manufacturers protocol for RNA isolation. This protocol resulted in approximately 5-6μg of RNA from each batch of 10 flies.

### Transcriptomic libraries

Poly-A mRNA was purified from total RNA using oligodT magnetic beads and strand-specific indexed libraries were prepared using the KAPA Stranded RNA-Seq kit followed by ten cycles of amplification using KAPA HiFi DNA polymerase (KAPA Biosystems). Libraries were quantified and pooled based on a post-PCR Agilent Bioanalyzer and 75 bp paired-end reads were generated on the Illumina HiSeq v4 following the manufacturer’s standard sequencing protocols. All raw sequencing reads are available (from the date of pending journal publication) on the European Nucleotide Archive under study accession number PRJEB35592.

### Read mapping and differential expression analysis

Reads were mapped to *P. papatasi* genome (Ppapl1 v1, Vectorbase^17^) using hisat2^52^. Reads which mapped uniquely and in their proper pair were extracted and used to assemble transcripts *de novo* with Cufflinks (Tuxedo suite)^53^. The newly assembled transcripts were combined with the VectorBase transcript assembly to create a new set of transcripts using CuffMerge. Both the sequences of the assembled transcripts and the new annotation file (.gtf) are included in the supplementary data files. Reads were then counted against the cufflinks-generated transcripts using featureCounts^54^. The counts data for the (two) technical replicates for each sample were collapsed prior importing into R for differential expression analysis (pairwise Wald tests) in DESeq2^55^.

## Acknowledgements

We thank the staff of the DNA pipelines at Wellcome Sanger Institute for sequencing and generating sequencing libraries. This work was supported by the European Commission, Horizon 2020 Infrastructure Infravec2 project (grant agreement No 731060, https://infravec2.eu). JS and PV were supported by ERD Funds, project CePaViP (CZ.02.1.01/16_019/0000759). MJS and JAC were supported by Wellcome via their core support for the Wellcome Sanger Institute (WSI) through grant 206194. Work in Oxford was supported by a Consolidator grant from the European Research Council (310912 Droso-Parasite, to PL), project grant BB/K003569 from the BBSRC (to PL) and a Wellcome Trust doctoral scholarship (to MAS).

## Supplementary figures

**Figure S1.**
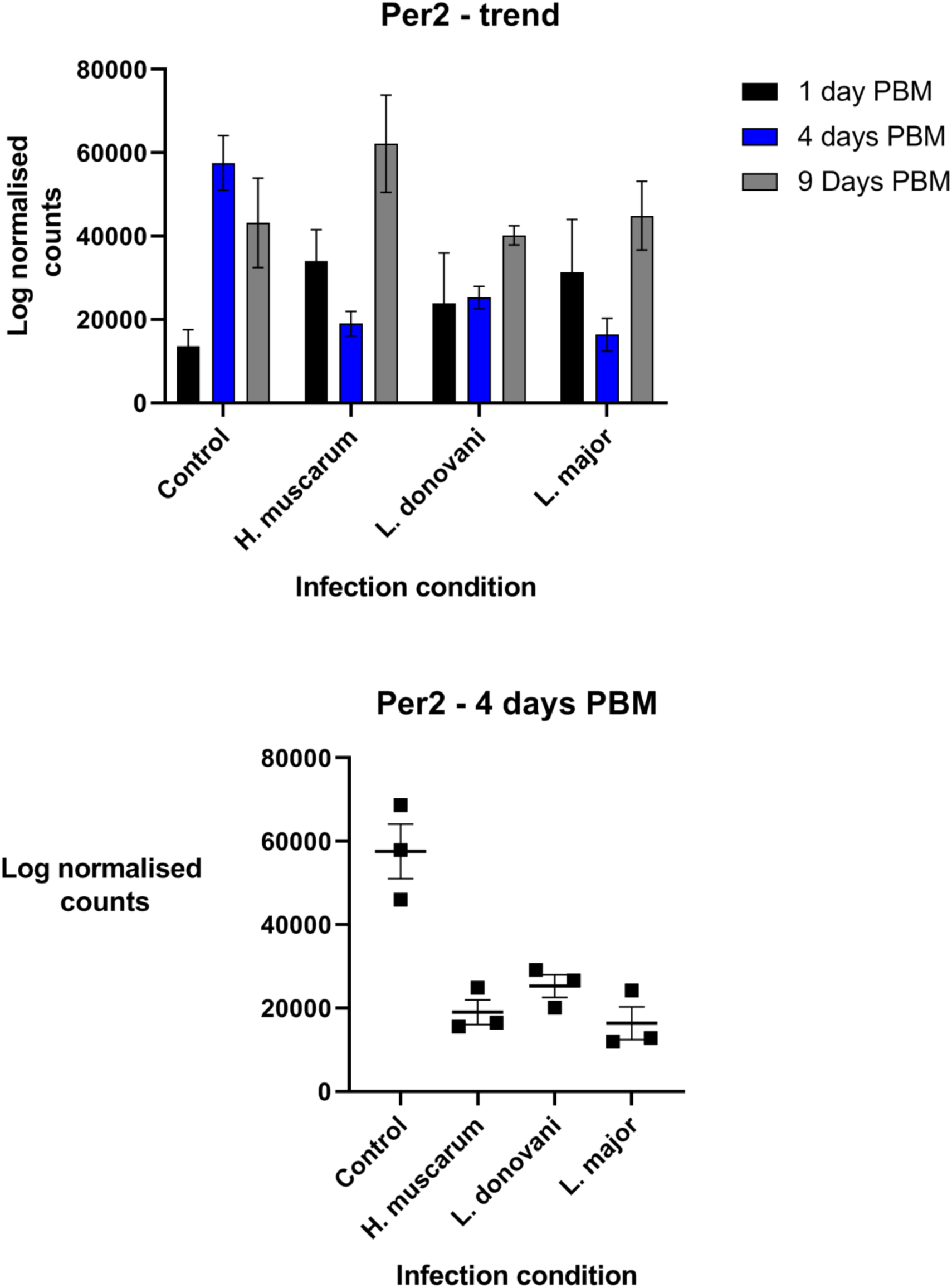
Log normalised transcript counts for Peritrophin 2 (Per2) in P. papatasi throughout infection. Error bars show the standard error of the mean.

## List of Supplementary Tables

**Table S1 – Read mapping summaries.** This table shows the read mapping information for each sample e.g. number of reads, percentage of read mapped etc.

**Table S2 – Transcripts associated with trypanosomatid presence in the blood meal.** This table shows the fold changes and differential regulation statistics (including p-values) for transcripts whose abundance differed between trypanosomatid fed flies and blood-fed control flies.

**Table S3 – Transcripts associated with specific trypanosomatids in the blood meal.** This table shows the fold changes and differential regulation statistics (including p-values) for transcripts whose abundance differed between trypanosomatid infections.

**Table S4 – Transcripts significantly differentially regulated between 1 day and 4 days post bloodmeal (blood-only) in *P. papatasi*.**

**Table S5 – Transcripts significantly differentially regulated between 4 day and 9 days post bloodmeal (blood-only) in *P. papatasi*.**

**Table S6 – Transcripts of interest which are differentially regulated between 1 day and 4 days post bloodmeal (blood-only) in *P. papatasi*.** This is a streamlined version of supplementary table 4 showing transcripts of interest discussed in the text.

**Table S7 – Transcripts of interest which are differentially regulated between 4 days and 9 days post bloodmeal (blood-only) in *P. papatasi*.** This is a streamlined version of supplementary table 4 showing transcripts of interest discussed in the text.

**Table S8 – Differential regulation statistics for transcripts of dipteran immune pathways of interest (Toll, Imd, DUOX and JAK-STAT) across samples. ns –** not significantly differentially regulated.

## List of supplementary data files

**Annotation file for *de novo* assembled transcripts merged with Ppal1 annotation (.gtf) Assembled transcript sequences (.fasta)**

